# Bacterial diversity along a 2600 km river continuum

**DOI:** 10.1101/010041

**Authors:** Domenico Savio, Lucas Sinclair, Umer Z. Ijaz, Philipp Stadler, Alfred P. Blaschke, Georg H. Reischer, Guenter Blöschl, Robert L. Mach, Alexander K.T. Kirschner, Andreas H. Farnleitner, Alexander Eiler

## Abstract

The bacterioplankton diversity in large rivers has thus far been undersampled, despite the importance of streams and rivers as components of continental landscapes. Here, we present a comprehensive dataset detailing the bacterioplankton diversity along the midstream of the Danube River and its tributaries. Using 16S rRNA-gene amplicon sequencing, our analysis revealed that bacterial richness and evenness gradually declined downriver in both the free-living and particle-associated bacterial communities. These shifts were also supported by beta diversity analysis, where the effects of tributaries were negligible in regards to the overall variation. In addition, the river was largely dominated by bacteria that are commonly observed in freshwaters. Dominated by the acI lineage, the freshwater SAR11 (LD12) and the *Polynucleobacter* group, typical freshwater taxa increased in proportion downriver and were accompanied by a decrease in soil and groundwater bacteria. Based on the River Continuum Concept, we explain these taxonomic patterns and the accompanying changes in alpha and beta diversity by the physical structure and chemical conditions coupled with the hydrologic cycle along the length of the river.

## Introduction

Streams and rivers link terrestrial and lentic systems with their marine counterparts and provide numerous essential ecosystem services. They supply drinking water, are used for irrigation, industry and hydropower, and serve as transport routes or for recreation. Of general importance is the role of lotic systems in biogeochemical nutrient cycling. Until recently, rivers and streams were mainly considered as pipes shuttling organic material and nutrients from the land to the ocean (Cole *et al.*, 2007). This view has begun to change as lotic and lentic systems are now considered more akin to “leaky funnels” in regard to the cycling of elements. Indeed, they play an important role in the temporary storage and transformation of terrestrial organic matter (Ensign and Doyle, 2006; Cole *et al.*, 2007; Withers and Jarvie, 2008; Battin *et al.*, 2009). As a result of recognising the role of rivers and streams in the carbon cycle (Richey *et al.*, 2002; Battin *et al.*, 2009; Raymond *et al.*, 2013), the study of the diverse, ongoing processes in the water column and sediments of lotic networks has received increasing interest (Kronvang *et al.*, 1999; Beaulieu *et al.*, 2010; Seitzinger *et al.*, 2010; Aufdenkampe *et al.*, 2011; Benstead and Leigh, 2012; Raymond *et al.*, 2013).

When attempting to model the mechanisms of nutrient processing in freshwater systems, bacteria are regarded as the main transformers of elemental nutrients and viewed as substantial contributors to the energy flow (Cotner and Biddanda, 2002; Battin *et al.*, 2009; Findlay, 2010; Madsen, 2011). However, in the case of open lotic systems such as rivers, there remains a lack of knowledge concerning the diversity of bacterial communities (Battin *et al.*, 2009). There is currently no agreement on the distinctness of river bacterioplankton communities from that of other freshwater systems or the variability of its diversity along entire rivers.

When summarising previous studies, it can be concluded that the abundant taxa comprising the riverine bacterioplankton resemble lake bacteria and can thus be regarded as “typical” freshwater bacteria (Zwart *et al.*, 2002; Lozupone and Knight, 2007; Newton *et al.*, 2011). In particular bacteria affiliated with the phyla of *Proteobacteria (*particularly *Betaproteobacteria), Actinobacteria*, *Bacteroidetes, Cyanobacteria a*nd *Verrucomicrobia* dominate the bacterial communities in rivers (Crump *et al.*, 1999; Zwart *et al.*, 2002; Cottrell *et al.*, 2005; Winter *et al.*, 2007; Lemke *et al.*, 2008; Mueller-Spitz *et al.*, 2009; Newton *et al.*, 2011; Liu *et al.*, 2012). A recent metagenome study corroborates a general dominance of the phyla *Proteobacteria* and *Actinobacteria,* and more specific the clear dominance of the cosmopolitan freshwater lineage acI of the phylum *Actinobacteria* in the Amazon river (Ghai *et al.*, 2011). The dominance of *Actinobacteria* and *Proteobacteria* in riverine bacterioplankton was also confirmed in two high-throughput sequencing studies on the Upper Mississippi River (USA; Staley *et al.*, 2013) and the Yenisei River (RUS; Kolmakova *et al.*, 2014). The former revealed a ubiquitous ‘core bacterial community’ to be present in the Upper Mississippi River (USA), whereas in the latter three distinctly different bacterial assemblages were identified based on beta-diversity analysis.

The longitudinal development of the bacterioplankton community along an entire river was so far only addressed in a study on the 354 km long River Thames (UK) (published during the review process of this manuscript; Read *et al.*, 2014). In this study, the authors observed a shift from a *Bacteroidetes*-dominated community in the headwaters to an *Actinobacteria* dominated community, leading them to conclude that bacterioplankton communities are formed by the process of succession. However, in the case of macroorganisms, the applicability of this concept to riverine communities has been challenged based on the argument that communities in each reach have a continuous heritage rather than an isolated temporal composition within a sequence of discrete successional stages (Vannote *et al.*, 1980).

For this reason, the River Continuum Concept (RCC; Vannote *et al.*, 1980) was proposed as the framework of choice to explain large-scale diversity patterns observed from headwater streams to large rivers. For macroorganisms, based on ‘diel temperature variability’, the RCC postulates that diversity increases from headwaters to medium-sized stream reaches, with a subsequent decrease towards the river mouth. Yet, there is much more to the RCC than this widely referred hump-shaped diversity pattern, as the RCC provides a comprehensive conceptual framework for the description of diversity patterns in large river systems. It does so i.a. by explicitly emphasizing that the physical structure coupled with the hydrological cycle form a templet for biological responses and result in consistent patterns of community structure and function (Vannote *et al.*, 1980). In particular, the RCC highlights the role of the ‘riparian zone’, ‘substrate’ availability, ‘flow’, and ‘food’ as important factors in determining community structure.

Here, we explain the diversity patterns of river bacterioplankton in the context of the RCC by utilising the results from a second-generation sequencing experiment detailing the bacterial community composition along a large river. Furthermore, we reveal how the variability in bacterioplankton diversity is related to the environmental variables along 2600 river kilometre from medium-sized reaches to the river mouth. We separately investigated the free-living communities and particle-associated communities by extracting two different size fractions (0.2-3.0 μm and >3.0 μm) for each sample. These two fractions have been shown to exhibit significant differences in activity and community dynamics in previous studies, justifying this distinction (Crump *et al.*, 1999; Velimirov *et al.*, 2011). The study site was the Danube River (Fig. 1), the second largest river in Europe by discharge and length. The Danube River drains a basin of approximately 801 000 km^2^; the area is populated with 83 million inhabitants and borders 19 countries (Sommerwerk *et al.*, 2010).

**Fig. 1.**
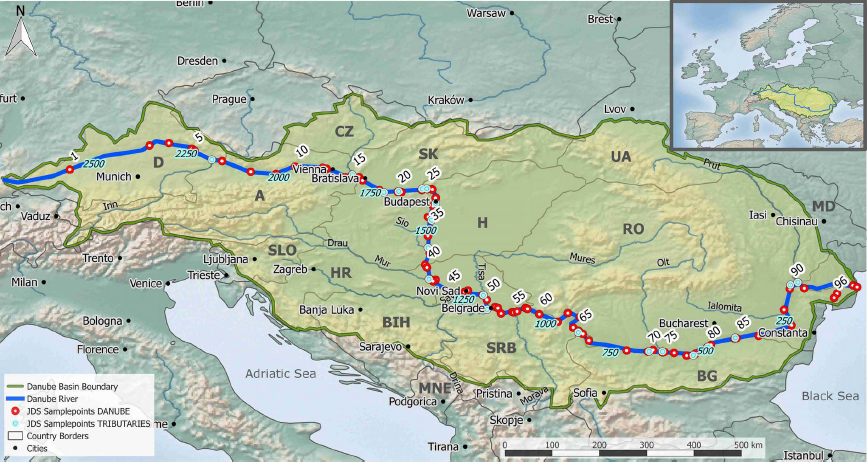
Overview- and detailed map of the Danube River catchment showing all sampling sites during the Joint Danube River Survey 2; red dots indicate sampling points in the midstream of the Danube River; blue dots represent sampling points in tributaries before merging in the Danube River. Blue-shaded font indicates official numbering of river starting with rkm 2600 at the source to rkm 0 at the river mouth. Country abbreviations and large cities are written in black. The map was created using Quantum GIS (Quantum GIS Development Team, 2011).

## Results

### Description of selected environmental parameter

In total, more than 280 individual parameters, including chemical, microbiological, ecotoxicological, radiological and biological parameters, were investigated within the Joint Danube Survey 2. Alkalinity, pH, nitrate concentration as well as concentration of dissolved silicates exhibited a gradually decreasing trend along the river as previously described by Liska and colleagues (2008) and illustrated in Fig. S2. Total phytoplankton biomass (Chl-a) showed a peak between river kilometre 1481 and 1107 (sites 38-55) with total bacterial production following a similar trend, whereas total suspended solid concentration increased considerably in the last 900 kilometres before reaching the Black Sea (Fig. S2).

### Core microbial community

In total, sequencing resulted in 1 572 361 sequence reads (further referred to as “reads”) after quality filtering, clustering into 8697 bacterial OTUs. The majority of bacteria-assigned OTUs (4402 out of 8697) were only represented by less than ten reads in the entire dataset. As a consequence, 3243 of 8697 OTUs (∽37%) were present in only one to four samples, and an additional 2219 OTUs (∽26%) were present in as few as five to nine samples. In addition to these rare OTUs, the core community of the Danube River, operationally defined by all OTUs that appeared in at least 90% of all samples, comprised 89 OTUs in the free-living bacterioplankton (0.2-3.0 μm) and 141 OTUs in the particle-associated microbes (>3.0 μm).

The cumulative contribution of OTUs based on their occurrence along the entire river is shown in Fig. 2A. for both analysed size fractions. On average, 81% of all reads of the free-living river community and 63% of all reads of the particle-associated river community were part of their respective core community. Based on this visualization, the core communities of both size fractions were defined by all OTUs that are present in 90% or more of all samples.

**Fig. 2.**
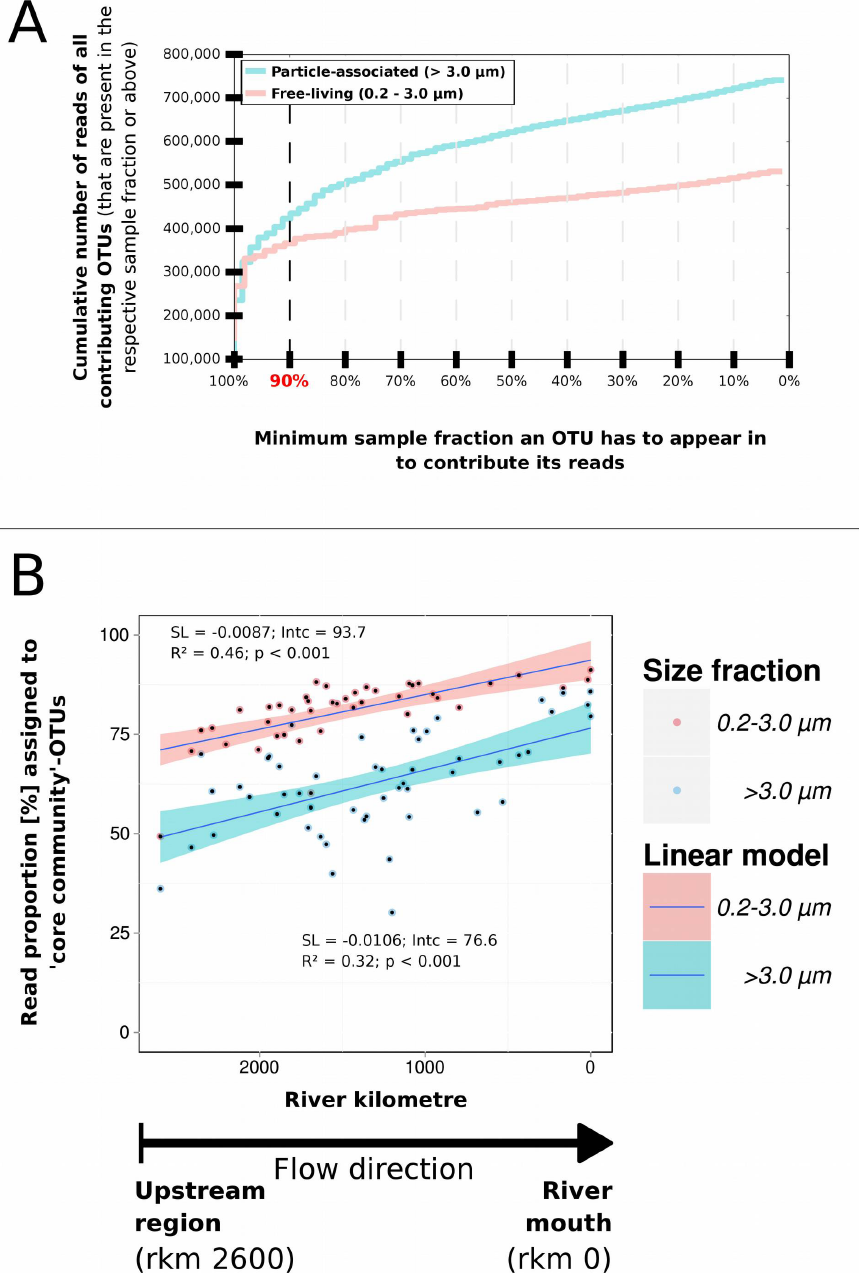
A. Cumulative graph of the absolute quantitative contribution of OTUs based on their occurrence in distinct fractions of samples. The X-axis displays the fraction of samples in % an OTU has to be present at the minimum to contribute its assigned reads; the Y-axis shows the cumulative number of reads corresponding to those OTUs that appear in the respective sample fraction or above. The blue line represents the particle-associated bacterial fraction (>3.0 μm); the red line shows the free-living bacterial fraction (0.2-3.0 μm). **B.** Gradual development of the read proportion assigned to the operationally defined “core communities” of the free-living and particle-associated fraction (all OTUs in the respective size fraction that occur in 90% of all river samples or above; compare Fig. 2A) along the Danube River from upstream (left; rkm 2600) to the river mouth (right; rkm 0). Red symbols indicate samples from the free-living fraction (0.2-3.0 μm); blue symbols indicate the particle-associated fraction (>3.0 μm); dark blue lines represent fitted linear models with confidence intervals of 0.95 in red and blue for the respective fractions. Detailed regression statistics are shown in the figure & Table 1.

**Table 1.** Summary of regression statistics (slope, intercept, multiple R-squared and p-value) for fitted linear models between Chao1 richness (Fig. 3A), Pielou’s evenness (J; Fig. 3B), core community proportions (Fig. 2B) and selected environmental parameter (Fig. S2) and river kilometre or mean discharge, respectively. FL: free-living community of the Danube River (0.2-3.0 μm); and PA: particle-associated Danube River community (>3.0 μm);

| **Correlations with River Kilomtre** | **Slope** | **Intercept** | **R<sup>2</sup>** | **p-value** |
| --- | --- | --- | --- | --- |
| Chao1 richness FL | 3.958E-01 | 6.305E+02 | -0.392 | <0.001 |
| Chao1 richness PA | 2.371E-01 | 1.721E+03 | -0.173 | <0.01 |
| Evenness FL | 4.226E-05 | 6.264E-01 | -0.483 | <0.001 |
| Evenness PA | 5.405E-05 | 6.973E-01 | -0.501 | <0.001 |
| Core community FL | -8.683E-03 | 9.374E+01 | 0.455 | <0.001 |
| Core community PA | -1.055E-02 | 7.664E+01 | 0.316 | <0.001 |
| Alkalinity | 3.671E-04 | 2.283E+00 | -0.423 | <0.001 |
| Nitrate | 4.384E-04 | 1.167E+00 | -0.512 | <0.001 |
| pH | 1.659E-04 | 7.575E+00 | -0.208 | <0.001 |
| Silicates dissolved | 1.112E-03 | 3.956E+00 | -0.430 | <0.001 |
| **Correlations with Mean Discharge** | **Slope** | **Intercept** | **R<sup>2</sup>** | **p-value** |
| Chao1 richness FL | -1.473E-01 | 1.649E+03 | -0.450 | <0.001 |
| Chao1 richness PA | -9.780E-02 | 2.372E+03 | -0.254 | <0.01 |
| Evenness FL | -1.519E-05 | 7.354E-01 | -0.598 | <0.001 |
| Evenness PA | -1.280E-05 | 8.177E-01 | -0.302 | <0.001 |

Fig. 2B shows the observed significant increase in relative quantitative contribution of the core communities in both fractions towards the river mouth. Regression analysis revealed similar slopes for the proportions of the free-living as well as particle-associated core-communities along the river, whereas the free-living fraction on average contributed a higher proportion (81%) to the whole community compared to the particle associated fraction (63%).

### Variability of diversity along the river

To follow patterns in alpha diversity, we calculated the Chao1 richness estimator and Pielou’s evenness index for both size fractions after rarefying all samples down to 7000 reads and discarding 36 samples with less reads. The estimated richness was consistently higher in the particle-associated fraction when compared to the free-living fraction (Wilcoxon rank sum test; p-value < 0.001) with averages of 2025 OTUs and 1248 OTUs, respectively. We observed the highest diversity of all samples in the upstream part of the Danube River, representing medium-sized stream reaches according to definitions of the RCC. Richness and evenness then gradually decreased downstream in both size fractions (Fig. 3A+B) as confirmed by the regression analysis using both river kilometre and mean discharge (Table 1). The comparison of the slopes for both size fractions revealed a steeper decline for estimated Chao1 richness in the free-living compared to the particle-associated fraction.

**Fig. 3.**
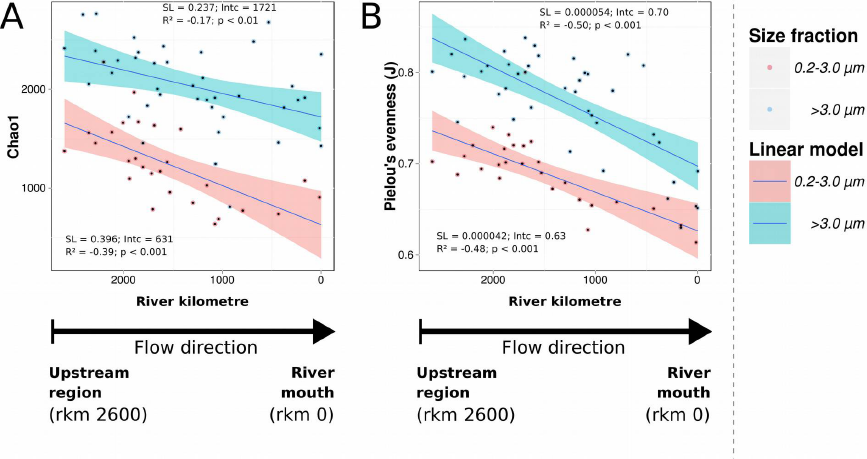
The gradual development of (**A**) the bacterial richness (Chao1) and (**B**) Pielou’s evenness (J) along the Danube River in the two size fractions, representing the bacterioplankton communities of 0.2-3.0 μm and >3.0 μm (corresponding to free-living and particle-associated bacterioplankton, respectively) from upstream (left; rkm 2600) to the river mouth (right; rkm 0). Red symbols indicate samples from the free-living fraction (n=27); blue symbols samples from the particle-associated fraction (n=40). Dark blue lines represent fitted linear models with confidence intervals of 0.95 in red and blue for the respective fractions. Detailed regression statistics are shown in the figure & Table 1.

To analyse variability in beta diversity, we first visualised the community changes along the continuum by applying non-metric multidimensional scaling (NMDS) to a Bray-Curtis dissimilarity matrix (Fig. 4). In both size fractions, we observed a significant relationship between community composition and river kilometre. While communities of both size fractions correlated significantly with pH, alkalinity, nitrate concentration and dissolved silicates, the particle-associated community additionally correlated with total bacterial production, phytoplankton biomass and total suspended solids (Table 2). For details, the dynamics of the correlating environmental parameters along the Danube River are shown in Fig. S2.

**Table 2.** Summary statistics of correspondence between environmental variables and the projections of bacterioplankton community samples in the NMDS ordination based on either free-living or particle-associated size fractions from the Danube River. The results were obtained using the function ‘envfit’ included in the R-package ‘vegan’ (Oksanen *et al.*, 2013).

| Variables | free-living<br>R<sup>2</sup> | particle-associated<br>R<sup>2</sup> |
| --- | --- | --- |
| River kilometre | 0.844*** | 0.795*** |
| Water temperature | 0.264** | 0.035 |
| Dissolved oxygen | 0.05 | 0.002 |
| pH | 0.330*** | 0.214** |
| Conductivity | 0.303* | 0.093 |
| Alkalinity | 0.626*** | 0.333*** |
| Ammonium | 0.117 | 0.004 |
| Nitrite | 0.285** | 0.254** |
| Nitrate | 0.705*** | 0.419*** |
| Organic nitrogen | 0.024 | 0.076 |
| Orthophosphate phosphorus | 0.186* | 0.099. |
| Total phosphorus | 0.069 | 0.059 |
| Silicates dissolved | 0.504*** | 0.515*** |
| Phytoplankton biomass (Chla) | 0.179* | 0.380*** |
| Total suspended solids | 0.150. | 0.345*** |
| Production total | 0.051. | 0.397*** |
| DOC | 0.042 | 0.183* |

Signif. codes: 0 '\*\*\*' 0.001 '\*\*' 0.01 '\*' 0.05 '.' 0.1 ' ' 1

**Fig. 4.**
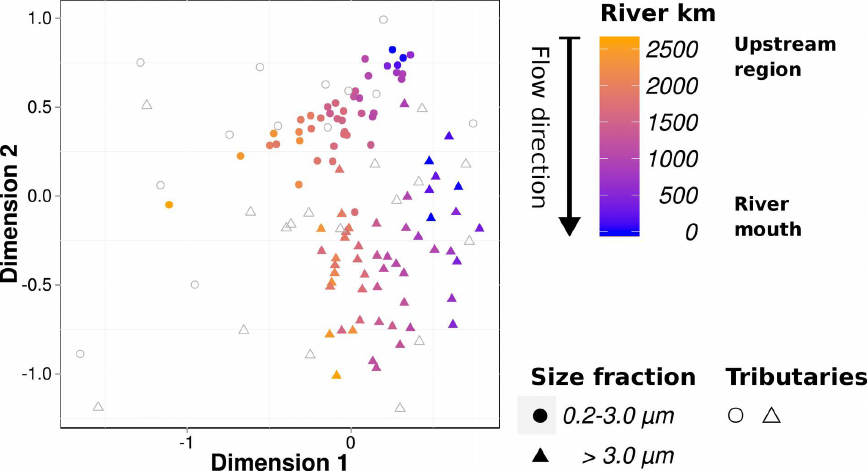
The visualisation of the beta diversity analysis based on the Bray-Curtis dissimilarity index shows the compositional dissimilarity between sites along the Danube River and its tributaries from upstream (left; rkm 2600) to the river mouth (right; rkm 0). The stress value of the non-metric multidimensional scaling (NMDS) was 0.17. Circles represent free-living bacterial communities (0.2-3.0 μm); triangles represent particle-associated bacterial communities (>3.0 μm). Open symbols display tributary samples, whereas full symbols indicate Danube River communities. The gradient from orange to blue via purple indicates the position of the sampling site upstream from the river mouth. The official assignment of river kilometres (rkm) for the Danube River is defined in a reverse fashion starting from the mouth (rkm 0) and progressing towards the source with our most upstream site (rkm 2600).

Other visual impressions from the NMDS are (1) that tributaries did not follow the general patterns and often formed outliers in the ordination space; (2) that there is a distinction in community composition between the two size fractions, which we confirmed by PERMANOVA analysis (R^2^=0.156, p-value<0.01); and (3) that there appears to be synchrony in the community changes of the two size fractions along the river’s course, which we statistically verified using a procrustes test (R=0.96, p<0.001). Furthermore, the application of a permutation test to the beta dispersion values of each size fraction revealed a higher variability in the >3.0 μm fraction when compared to the 0.2-3.0 μm fraction (p-value=0.002) (see Fig. S3).

### Typical river bacterioplankton

Along the river, the bacterioplankton community was dominated by *Actinobacteria*, *Proteobacteria*, *Bacteroidetes*, *Verrucomicrobia* and candidate division OD1, with an increase of reads assigned to the phylum *Actinobacteria* in the free-living size fraction downriver (Fig. S1). On the contrary, *Bacteroidetes*-assigned reads decreased significantly in the free-living fraction, whereas in the particle-associated fraction, these trends in phylum composition were less pronounced. In addition to assigning reads to the phylum level, we taxonomically annotated 9322 OTUs using similarity searches against the database of freshwater bacteria 16S rRNA sequences developed by Newton and colleagues (2011). The analysis revealed that up to 80% of the free-living and more than 65% of the particle-associated bacterial population inhabiting the Danube could be assigned to previously described freshwater taxa (Fig. 5B). In particular, these included representatives of the LD12-tribe belonging to the subphylum of *Alphaproteobacteria*, as well as the acI-B1-, acI-A7- and acI-C2*-*tribes belonging to the phylum *Actinobacteria*.

**Fig. 5.**
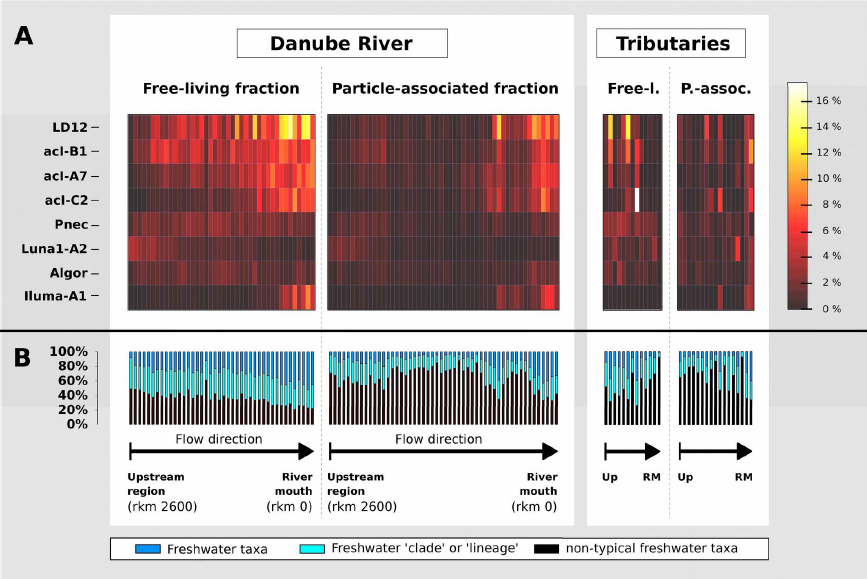
A heat map (**A**) revealing the dynamics of the eight most abundant typical freshwater tribes along the Danube River according to Newton *et al.*, 2011. The gradient from black red via yellow to white indicates the relative quantitative contribution of the respective tribe to all sequence reads in any one sample, with a maximum of 16%. Panel (**B**) shows the overall contribution of typical freshwater tribes, clades and lineages (Newton *et al.*, 2011) to the river bacterioplankton amplicon sequences along the river; Black bars represent reads that could not be matched to sequences of the used freshwater database (Newton *et al.*, 2011) neither on tribe-, clade-or lineage-level (named ‘non-typical freshwater taxa’). ‘Freshwater taxa’ and ‘Freshwater clade or lineage’ represent all reads that could be matched to sequences of the used freshwater database at the respective similarity-level. Samples of the Danube River as well as the investigated tributaries are arranged from left to the right, with increasing distance from the source and separated for the respective size fractions.

Interestingly, in the free-living size fraction, we observed a clear increase in the relative abundance of the four above mentioned tribes towards the river mouth (Fig. 5A), contributing up to 35% of the community. This increase in the contribution of these four tribes was accompanied by a general increase of the relative quantitative contribution of OTUs matching other freshwater tribes, lineages or clades according to Newton and colleagues (2011) as visualised in Fig. 5B. On the contrary, the number of OTUs not matching any sequence of the freshwater database either at tribe-, clade-or at lineage-level was decreasing (Fig. 5B, labelled “non-typical freshwater taxa”), raising the suspicion that these OTUs originated from non-aquatic sources. Particularly in the particle-associated fraction, typical freshwater taxa were less common (Fig. 5B).

To confirm the non-aquatic origin of certain OTUs, we first blasted a representative for each of the 8697 bacterial OTUs against the NCBI-NT database; next, any environmental descriptive terms occurring in the search results were retrieved and classified according to the Environmental Ontology (EnvO; Buttigieg *et al.*, 2013) terminology. PERMANOVA analysis of the EnvO-classified data revealed a significant difference in variance between the two size fractions (PERMANOVA; R^2^=0.42, p<0.0001). Restricting the analysis to particular EnvO terms such as ‘groundwater’ and ‘soil’ terms suggests that the proportion of bacteria potentially originating from these two sources decreased towards the river mouth (Fig. 6A and B), while ‘river’ and ‘sediment’ terms did not follow either a downriver or upriver trend. In particular, we could only identify four OTUs receiving an EnvO dominated by the ‘river’ term.

**Fig. 6.**
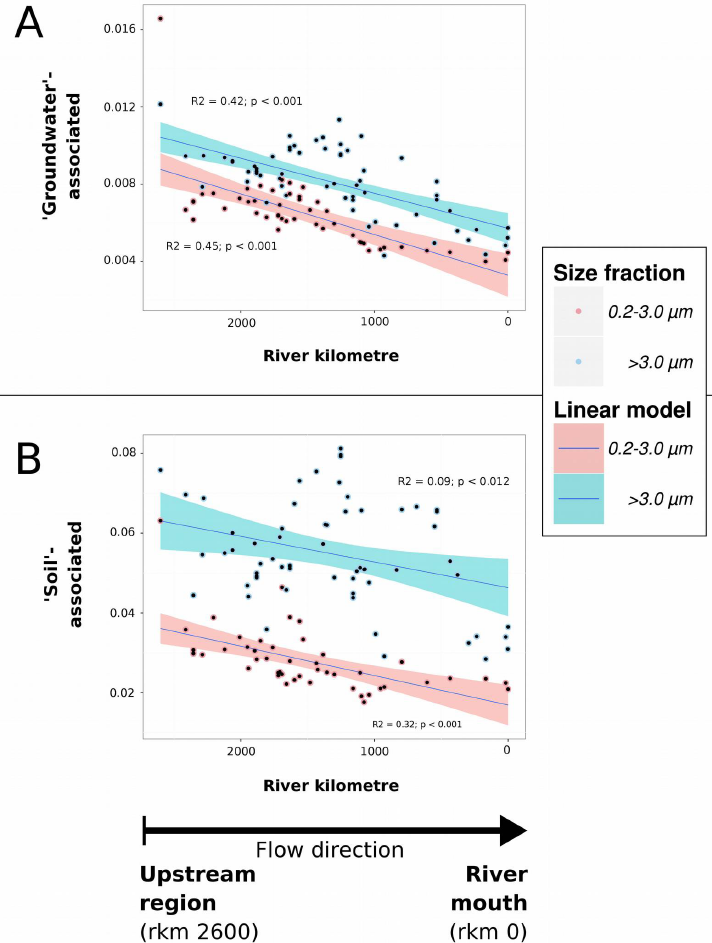
Results from the SEQenv analyses scoring sequences according to their environmental context. The Y-axis represents the proportion of (**A**) ‘groundwater’ and (**B**) ‘soil’ terms associated with sequence reads per sample along the Danube River (X-axis). Red symbols indicate samples from the 0.2-3.0 μm fraction (n=27), and blue symbols indicate samples from the >3.0 μm fraction (n=40). Dark blue lines represent fitted linear models with confidence intervals of 0.95 in red and blue for the respective fractions. Detailed regression statistics are given in the figure.

As with every homology based assignments, SeqEnv results are affected by database entries; for example the under-representation of entries from rivers compared to lakes likely discriminates against the ‘river’ term. Thus, a larger number of typical river bacterioplankton may exist than detected by our analysis.

## Discussion

### Explaining patterns of bacterioplankton diversity in the framework of the River Continuum Concept

The tremendous diversity within the microbial communities inhabiting all types of (aquatic) environments is being revealed by a rapidly increasing number of studies applying high-throughput sequencing technologies (e.g. Sogin *et al.*, 2006; Andersson *et al.*, 2009; Galand *et al.*, 2009; Eiler *et al.*, 2012; Peura *et al.*, 2012). At the same time, many mechanisms modulating this diversity have been suggested including ‘mass effect’, dispersal limitations and environmental condition based sorting (‘species sorting’) which vary widely in importance depending on the environment (Leibold *et al.*, 2004; Besemer *et al.*, 2012; Hanson *et al.*, 2012; Lindström and Langenheder, 2012; Szekely *et al.*, 2013).

Combining ours and previous results (Besemer *et al.*, 2012, 2013; Crump *et al.*, 2012; Staley *et al.*, 2013; Read *et al.*, 2014), we propose that the bacterioplankton diversity in a large river network is expected to be highest in headwaters, and from thereon decreases towards river mouths. This pattern of decreasing diversity from source to “sink”, we argue, can be explained within the framework of the RCC (Vannote *et al.*, 1980) by considering the underlined factors like ‘riparian influence’, ‘substrate’, ‘flow’ and ‘food’.

Regarding bacterioplankton, we propose that particularly the ‘riparian influence’ gains crucial importance in shaping a divergent diversity pattern compared to macroorganisms. The disparity, we suggest, has following reasons: (i) The primarily passive transport of bacterioplankton contrasts the habitat-restriction of macroorganisms like aquatic invertebrates, fish or macrophytes, which is based on their motility or sessility; (ii) as such the large contact zone of small headwaters with the surrounding environment (soil and groundwater) can constantly contribute allochthonous bacteria to the river community (Besemer *et al.*, 2012; Crump *et al.*, 2012); (iii) these allochthonous microbes, imported from soil and groundwater communities, harbour a much higher diversity when compared to planktonic communities (e.g., Crump *et al.*, 2012); and (iv) should be at least temporarily capable of proliferating in their new lotic environment, which makes them constitutive members of the community when compared to, e.g., terrestrial insects that fall or are washed into streams or rivers.

Besides the high impact from the riparian zone on headwater bacterioplankton communities, previously also merging tributaries or microbial pollution sources have been argued to potentially affect the river communities by providing allochthonous particles and bacteria. In this regard, our results of a gradual community shift and a linear increase of the core communities’ relative abundance in both size fractions (Fig. 2A+B) contrast this view. This tributary-independent development of the midstream community is furthermore supported by the rapidly decreasing number of ‘first-time occurrences’ of OTUs from upstream to downstream (Fig. S4). As an explanation, we provide the long mixing times of the incoming water and the restrained dilution that this entails as it was previously suspected by Velimirov and colleagues (2011). Later, Kolmakova and colleagues (2014) reported the phenomenon of a parallel flow of water from tributaries also for the receiving Yenisei river.

Besides the dominant role of allochthonous bacteria in headwaters (‘mass effects’) and mostly negligible impacts from tributaries to the midstream communities’ composition, the factors shaping bacterial diversity downriver are physical (‘substrate’) and chemical (‘food’ in the form of dissolved organic matter) environmental conditions. Resulting ‘species sorting’ is supported by the observed simultaneous decrease in evenness together with bacterial richness in both size fractions. A comparable rise of few and more competitive species was already implied in the RCC for macroorganisms from medium-sized reaches towards river mouths (Vannote *et al.*, 1980). These dynamics do not only imply an increase in competition, but also in the rate of local extinction (Leibold *et al.*, 2004; Crump *et al.*, 2012).

One specific environmental factor often involved in bacterial competition is the concentration and quality of dissolved organic matter (Eiler *et al.*, 2003; Fierer *et al.*, 2007). In this regard, the RCC proposes that labile allochthonous organic compounds are rapidly used at upstream sites where the stream has its maximum interface with the landscape. Meanwhile, more refractory and relatively high molecular weight compounds are thought to be exported downstream and to accumulate along the river (Vannote *et al.*, 1980). This supports our hypothesis of a competitive advantage of downstream-dominant OTUs in utilising increasingly available, nutrient-poor organic compounds, as reflected by the increasing relative abundance of typical freshwater taxa such as LD12 and acI. These taxa represent small cells with an oligotrophic lifestyle (Salcher *et al.*, 2011; Garcia *et al.*, 2013), thus complying with the observation of a general trend towards smaller cells along the Danube River (Velimirov *et al.*, 2011) and coinciding with decreasing concentrations of nutrients and dissolved organic matter.

Nevertheless, to prove the role of dissolved and solid organic matter sources in the apparent decline of richness towards the river mouth, an assessment of the organic matter composition as well as its bioavailability has to be included in future studies. Additionally, loss factors (such as sedimentation) and selective top-down control (such as grazing and viral lysis) have been shown to vary over environmental gradients and to substantially influence microbial diversity (Ayo *et al.*, 2001; Langenheder and Jürgens, 2001; Weinbauer, 2004; Pernthaler, 2005; Bouvier and Del Giorgio, 2007).

### Particles as hotspots of diversity in river bacterioplankton

Besides a linear decrease in richness and evenness (α-diversity) in both size fractions along the river, we observed a consistently higher richness in the particle-associated communities when compared to those of the free-living fraction. Similar observations were reported in studies on (coastal) marine environments as well as lentic freshwater environments (Bižić-Ionescu *et al.*, 2014; Mohit *et al.*, 2014). Bižić-Ionescu and colleagues ascribed the higher α-diversity in the particle-associated community to the high heterogeneity in the particle microenvironment. Similarly, we suggest that the higher richness is the result of elevated availability of distinct ecological niches inside and on the surface of particles when compared to the surrounding water column. Furthermore, a large spectrum of niches is given by the high heterogeneity amongst particles which should include mobilised sediments, living organisms such as planktonic algae or zooplankton and detritus derived from terrestrial and aquatic sources. The presence of diversely colonised particles of different age, origin and composition based on the observation of a higher richness in the particle-associated fraction was also recently suggested by Bižić-Ionescu *et al.* (2014). The variability in particle age and origin is also supported by our results of the compositional changes as well as the variations in EnvO terms attributed to the particle-associated bacterial communities.

### Towards a typical freshwater bacteria community along the river

Focusing on the taxonomic composition, our data shows that so-called “typical” freshwater bacteria, including members of the acI lineage (c.f. Newton *et al.*, 2011), the freshwater SAR11 group (LD12) and the *Polynucleobacter* genus, formed a major part of the bacterial “core community”, particularly in the free-living fraction. The observation that very few taxa dominate the riverine bacterioplankton communities is also consistent with findings of Staley and colleagues (2013), where the core community-OTUs were primarily assigned to phyla that were often reported to be highly abundant in river systems (*Proteobacteria, Actinobacteria, Bacteroidetes, Cyanobacteria or Verrucomicrobia*). Additionally, the dominance of typical freshwater taxa often observed in lakes corroborates the fact that river bacterioplankton resembles those of lakes (Zwart *et al.*, 2002; Lozupone and Knight, 2007; Newton *et al.*, 2011).

On a higher taxonomic level, we observed an increase in the quantitative contribution of the phylum *Actinobacteria* (including lineage acI) accompanied by a decreasing contribution of *Bacteroidetes-*assigned reads along the river. A very similar observation was reported recently in a study on the unfractionated bacterioplankton community along the 354 kilometre long Thames River (UK), leading the authors to the suggestion that community structure is shaped by the process of succession (Read *et al.*, 2014).

However, ecological succession is a phenomenon or process by which a community undergoes non-stochastic changes following a disturbance or initial colonization of a new habitat, which was argued not to be the case when considering the flow of water along a river (Vannote *et al.*, 1980). In particular when considering the influence of the riparian zone and the accompanied inputs of allochthonous bacteria as important factors in determining patterns in bacterioplankton diversity, the RCC becomes a much better suited framework for the description of patterns in bacterioplankton diversity. Furthermore, the RCC provides a framework for incorporating internal processes like ‘species sorting’ and its varying importance along a river network. However, although we were able to show that the contribution of ‘mass-effects’ and ‘species sorting’ in determining community composition is linked to distance from the river-source, the links between the patterns of diversity and ecosystem function remain to be revealed in large rivers.

## Experimental Procedures

### Supporting data

Within the frame of the Joint Danube Survey 2, a wide range of chemical and biological parameters was collected (Liska *et al.*, 2008). All data, sampling methods as well as analytical methods are made publicly available via the official website of the International Commission for the Protection of the Danube River (ICPDR;http://www.icpdr.org/wq-db/). Selected data from JDS 1 & 2 were published previously in several studies (Kirschner *et al.*, 2009; Janauer *et al.*, 2010; Velimirov *et al.*, 2011; von der Ohe *et al.*, 2011).

### Study sites and sample collection

Samples were collected within the frame of the second Joint Danube Survey project (JDS 2) in 2007. The overall purpose of the Joint Danube Surveys is to produce a comprehensive evaluation of the chemical and ecological status of the entire Danube River on the basis of the European Union Water Framework Directive (WFD) (Liska *et al.*, 2008). During sampling from Aug 15th to Sept 26th 2007, 75 sites were sampled along the mainstream of the Danube River along its shippable way from river kilometre (rkm) 2600 to the river mouth at rkm 0 (Kirschner *et al.*, 2009) as shown in Fig. 1. In addition, 21 samples from the Danube’s major tributaries and branches were included. At the most upstream sites, the Danube River is representative of a typical stream of the rithron and characterised by its tributaries Iller, Lech and Isar (Kavka and Poetsch, 2002). The trip took 43 days and is equivalent to the average retention time of a water body in this part of the Danube River (for discussion of this issue, see Velimirov *et al.*, 2011). Samples were collected with sterile 1 L glass flasks from a water depth of approximately 30 cm. Glass flasks were sterilised by rinsing with 0.5% HNO3 and autoclaving them. For DNA extraction of the particle-associated bacterioplankton depending on the biomass concentration, 120-300 mL river water was filtered through 3.0 μm pore-sized polycarbonate filters (Cyclopore, Whatman, Germany) by vacuum filtration. The filtrate, which represented the bacterioplankton fraction smaller than 3.0 μm (later referred to as “free-living” bacterioplankton), was collected in a sterile glass bottle and subsequently filtered through 0.2 μm pore-sized polycarbonate filters (Cyclopore, Whatman, Germany). The filters were stored at -80 °C until DNA extraction.

### DNA extraction and quantification of bacterial DNA using quantitative PCR (qPCR)

Genomic DNA was extracted using a slightly modified protocol of a previously published phenol-chloroform, bead-beating procedure (Griffiths *et al.*, 2000) using isopropanol instead of polyethylene glycol for DNA precipitation. Total DNA concentration was assessed applying the Quant-iT™ PicoGreen® dsDNA Assay Kit (Life Technologies Corporation, USA), and 16S rRNA genes were quantified using domain-specific quantitative PCR. Quantitative PCR reactions contained 2.5 μL of 1:4 and 1:16 diluted DNA extract as the template, 0.2 μM of primers 8F and 338 (Frank *et al.*, 2007; Fierer *et al.*, 2008) targeting the V1-V2 region of most bacterial 16S rRNA genes and iQ™ SYBR® Green Supermix (Bio-Rad Laboratories, Hercules, USA). All primer information is available in Table S1. The ratios of measured 16S rRNA gene copy numbers in the different sample dilutions that deviated markedly from 1 after multiplication with the respective dilution factor were interpreted as an indicator for PCR-inhibition.

### Preparation of 16S rRNA gene amplicon libraries

For the preparation of amplicon libraries, 16S rRNA genes were amplified and barcoded in a two-step procedure to reduce PCR bias that is introduced by long primers and sequencing adaptor-overhangs (Berry *et al.*, 2011). We followed the protocol as described by Sinclair *et al.* (unpublished, see Supporting information). In short, 16S rRNA gene fragments of most bacteria were amplified by applying primers Bakt_341F and Bakt_805R (Herlemann *et al.*, 2011; Table S1) targeting the V3-V4 variable regions. In 25 μL reactions containing 0.5 μM primer Bakt_341F and Bakt_805R, 0.2 μM dNTPs (Invitrogen), 0.5 U Q5 HF DNA polymerase and the provided buffer (New England Biolabs, USA), genomic DNA was amplified in duplicate in 20 cycles. To use equal amounts of bacterial template DNA to increase the comparability and reduction of PCR bias, the final volume of environmental DNA extract used for each sample was calculated based on 16S rRNA gene copy concentration in the respective sample determined earlier by quantitative PCR (see above). For 105 samples, the self-defined optimum volume of environmental DNA extract corresponding to 6.4 × 10^5^ 16S rRNA genes was spiked into the first step PCR reactions; however, for 27 samples, lower concentrations were used due to limited amounts of bacterial genomic DNA or PCR inhibition detected by quantitative PCR (see above). These 132 samples included eight biological replicates. Prior to the analysis, we removed four samples due to their extremely low genomic DNA concentrations and 16S rRNA gene copy numbers. Duplicates of PCR products were pooled, diluted to 1:100 and used as templates in the subsequent barcoding PCR. In this PCR, diluted 16S rRNA gene amplicons were amplified using 50 primer pairs with unique barcode pairs (Sinclair *et al.*, in review; Table S1). The barcoding PCRs for most samples were conducted in triplicates analogous to the first PCR (n=100). The remaining 32 samples that had weak bands in first step PCR due to low genomic template DNA concentrations or high sample dilution were amplified in 6-9 replicates to increase amplicon DNA yield. Barcoded PCR amplicons were pooled in an equimolar fashion after purification using the Agencourt AMPure XP purification system (Beckman Coulter, Danvers, MA, USA) and quantification of amplicon-concentration using the Quant-iT^(tm)^ PicoGreen® dsDNA Assay Kit (Life Technologies Corporation, USA). Finally, a total of 137 samples including 5 negative controls resulted in four pools for sequencing.

### Illumina^®^ sequencing

The sequencing was performed on an Illumina^®^ MiSeq at the SciLifeLab SNP/SEQ sequencing facility hosted by Uppsala University. For each pool, the library preparation was performed separately following the TruSeq Sample Preparation Kit V2 protocol (EUC 15026489 Rev C, Illumina) with the exception of the initial fragmentation and size selection procedures. This involves the binding of the standard sequencing adapters in combination with separate Illumina^®^-specific MID barcodes that enables the combination of different pools on the same sequencing run (Sinclair *et al.*, in review). After pooling, random PhiX DNA was added (5%) to provide calibration and help with the cluster generation on the MiSeq’s flow cell.

### 16S rRNA gene amplicon data analysis

The sequence data were processed as outlined in Sinclair *et al.* (in review). In short, after sequencing the libraries of 16S rRNA amplicons, the read pairs were demultiplexed and joined using the PANDAseq software v2.4 (Masella *et al.*, 2012). Next, reads that did not bear the correct primer sequences at the start and end of their sequences were discarded. Reads were then filtered based on their PHRED scores. Chimera removal and OTU (operational taxonomic unit) clustering at 3% sequence dissimilarity was performed by pooling all reads from all samples together and applying the UPARSE algorithm v7.0.1001 (Edgar, 2013). Here, any OTU containing less than two reads was discarded. Each OTU was subsequently taxonomically classified by operating a similarity search against the SILVAmod database and employing the CREST assignment algorithm (Lanzén *et al.*, 2012). Plastid, mitochondrial and archaeal OTUs were removed. In addition, OTUs were also taxonomically annotated against the freshwater database (Newton *et al.*, 2011) using the same method. If necessary, OTU rarefying for the purpose of standardising sequence numbers was performed using the ‘rrarefy’-function implemented in the R-package vegan (Oksanen *et al.*, 2013). For alpha-diversity analysis (Chao1-richness estimator and Pielou’s evenness), we rarefied down to 7000 and 2347 reads per sample, respectively. This was based on one study revealing that for water samples a sequencing depth 5000 16S rRNA gene reads per sample captured more than 80% of the trends in Chao1-richness and Pielou’s evenness (Lundin *et al.*, 2012). Furthermore, this study could show that for water samples 1000 reads per sample explained to 90% the trends in beta-diversity (Bray-Curtin dissimilarity index. By rarefying down to 2347, which was the read number of the sample with the lowest reads, all samples could be included in the beta-diversity analysis. Diversity measures, statistical analyses and plot-generation were conducted in R (R Core Team, 2013) using python scripts. The habitat index for the top 5000 OTUs was determined using the SEQenv pipeline (http://environments.hcmr.gr/seqenv.html). The SEQenv pipeline retrieves hits to highly similar sequences from public repositories (NCBI Genbank) and uses a text mining module to identify Environmental Ontology (EnvO; Buttigieg *et al.*, 2013) terms mentioned in the associated contextual information records (“Isolation Source” field entry for genomes in Genbank or associated PubMed abstracts). At the time of running SEQenv on our dataset (version 0.8), there were approximately 1200 EnvO terms organised into three main branches (namely, *environmental material*, *environmental feature*, and *biome*). However, we used SEQenv to retrieve a subset of these terms, i.e., those that contain “Habitat” (ENVO:00002036). Raw sequence data were submitted to the NCBI Sequence Read Archive (SRA) under accession number SRP045083.

### General description of sequences

In total, DNA was extracted and sequenced from 132 filtered water samples originating from the Danube River and its tributaries. In addition, the same procedure was applied to 5 negative control samples. The sequencing yielded 2 030 029 read pairs ranging from 3451 to 24 873 per sample. After quality filtering and mate-pair joining as outlined in Sinclair *et al.* (in review; see Supporting information), 1 572 361 sequence reads (further referred to as “reads”) were obtained. The OTU clustering resulted in 8697 OTUs after the removal of all Plastid-, Mitochondrion-, *Thaumarchaeota*-, *Crenarchaeota*- and *Euryarchaeota*-assigned OTUs. Archaea-assigned OTUs were removed because of the use of bacteria-specific primers not giving a representative picture of the targeted Archaea-community. The undesirable Plastid-, Mitochondrion- and Archaea-sequences represented 19.1% of the reads and accounted for 625 OTUs. Next, for the alpha diversity analysis, we excluded any sample with less than 7000 reads, resulting in 8241 OTUs in the remaining 88 samples. By contrast, for the beta diversity analysis, which is less affected by rare OTUs, all samples were randomly rarefied to the lowest number of reads in any one sample in order to include a maximum number of samples in the analysis. This brought every sample down to 2347 reads, and any OTU containing less than two reads was discarded, which brought the total OTU count to 5082.

## Acknowledgements

This study was supported by the Austrian Science Fund (FWF) as part of the DKplus “Vienna Doctoral Program on Water Resource Systems” (W1219-N22) and the FWF project P25817-B22, as well as the research project “Groundwater Resource Systems Vienna” in cooperation with Vienna Water (MA31). AE and LS are funded by the Swedish Foundation for Strategic Research (ICA10-0015). Infrastructure (cruise ships, floating laboratory) and logistics for collecting, storing and transporting samples were provided by the International Commission for the Protection of the Danube River (ICPDR). The analyses were performed using resources provided by the SNIC through the Uppsala Multidisciplinary Center for Advanced Computational Science (UPPMAX) under project “b2011035”.

### Conflict of Interest Statement

The authors declare no conflict of interest.

## Supporting information

**Fig. S1.**
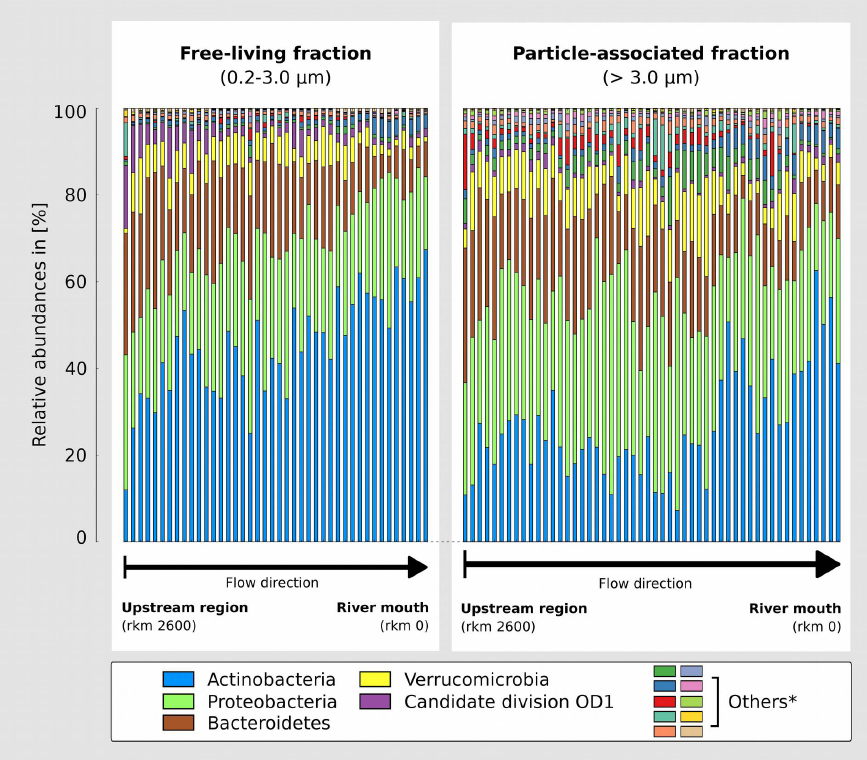
Phylum-level taxonomic composition of the bacterial communities along the Danube River. The Y-axis shows the read proportions assigned to the five most abundant phyla in the free-living fraction (left) and the particle-associated fraction (right). Lower abundant phyla were summarised to the fraction ‘Others’ due to their low proportions. Samples are arranged from left to the right representing sequence from upstream (rkm 2600) to river mouth at the Black Sea (rkm 0).

**Fig. S2.**
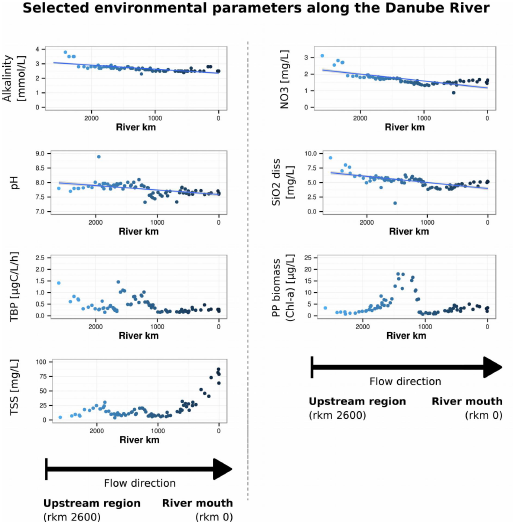
Development of selected environmental parameters along the Danube River from the upstream region (rkm 2600; left) to the river mouth at the Black Sea (rkm 0; right). Left panel: Alkalinity, pH, Total bacterial production (TBP), Total suspended solids (TSS); Right panel: Nitrate (NO_3_^-^), dissolved silicates (SiO_2_ diss) and Phytoplankton biomass (Chl-a) [PP biomass (Chl-a)]. Regression statistics of fitted linear models are given in Table 1 for the parameters Alkalinity, pH, NO_3_^-^, and SiO_2_ diss.

**Fig. S3.**
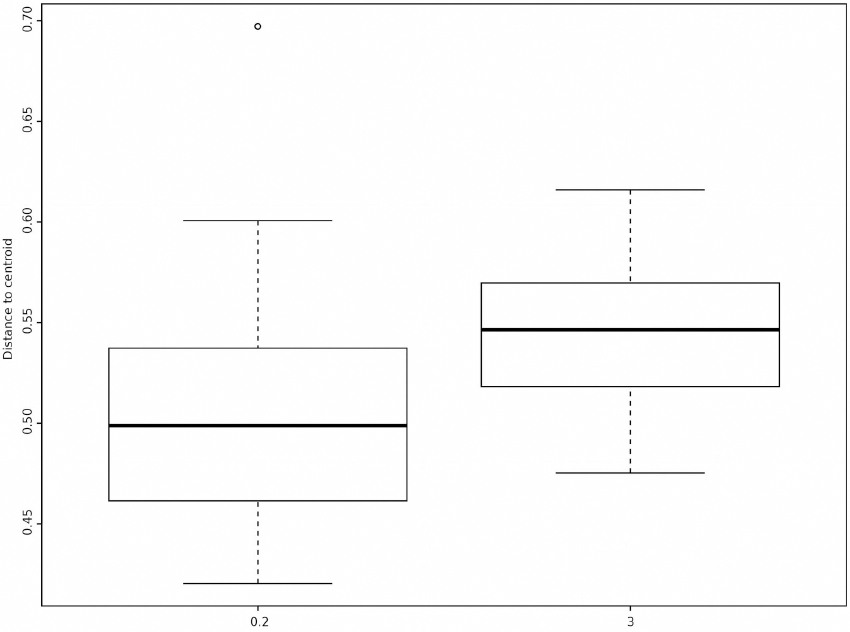
Boxplot of variability in bacterial communities in different size fractions (0.2-3.0 μm and >3.0 μm) based on betadispersion of Bray-Curtis dissimilarities. Left: Variability (distance from centroid) in the free-living bacterial community; Right: Variability in the attached bacterial community.

**Fig. S4.**
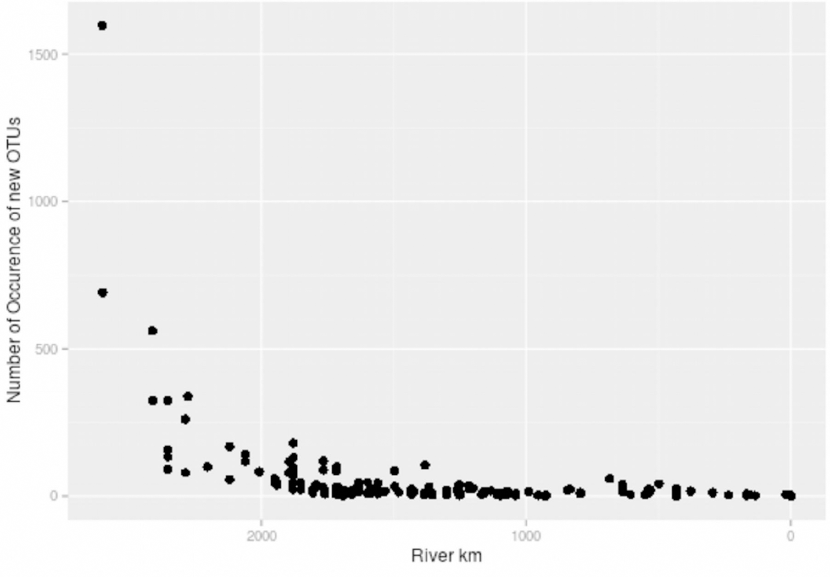
First occurrence plot of OTUs along the Danube River. Plotted are the numbers of OTUs occurring for the first time at the respective river kilometre (rkm) of each sampling sites.

